# Brainstem modulation of nociception by periaqueductal gray neurons expressing the mu-opioid receptor

**DOI:** 10.1101/2022.08.12.503787

**Authors:** Eileen Nguyen, Michael C. Chiang, Catherine Nguyen, Sarah E. Ross

## Abstract

Pharmacological manipulations directed at the periaqueductal gray (PAG) have revealed the importance of the mu-opioid receptor in the endogenous pain-modulatory system. Despite the clear role for opioidergic signaling within the PAG for the descending modulation of pain, the molecular and anatomical characterizations of neurons containing the mu-opioid receptor remain elusive. Using combinatorial anatomical, optogenetic, and chemogenetic approaches, we delineate a supraspinal pathway centered on PAG^MOR^ neurons in the modulation of pain and itch behaviors. We found that chemogenetic manipulations of PAG^MOR^ neurons in assays of nociception unveiled complex results; whereas activation of these neurons generally facilitated responses to noxious stimuli and jumping behaviors on the hotplate assay, opposing patterns were observed with reflexive responses to sensory testing. Activation of PAG^MOR^ neurons also robustly inhibited itch. These dichotomous findings across distinct types of sensory testing emphasize the contextual behavioral expression of nociception using reflexive and noxious paradigms. Lastly, we uncovered the role for PBN projections in the PAG that modulate pain in an uninjured, post-surgical state of latent sensitization.

## Introduction

Supraspinal structures including the midbrain periaqueductal gray (PAG) represent highly organized anatomical systems for the inhibition and facilitation of pain (Behbehani and Fields, 1979; Moreau and Fields, 1986; Budai et al., 1998). The PAG has been shown to modulate pain behaviors through a descending pathway involving the rostral ventromedial medulla (RVM) and spinal cord and is also a major component of the endogenous opioid analgesic system (Basbaum and Fields, 1978).

Pharmacological manipulations of the PAG have provided critical insight into the types of receptors that participate in the descending modulation of pain. For example, the injection of a mu-opioid receptor (MOR) agonist, such as DAMGO or morphine, into the PAG results in elevation of sensory thresholds and analgesia (Lewis and Gebhart, 1977; Carstens et al., 1990; Yaksh, 1997; Loyd et al., 2008b, 2008a). Local microinjections of the mu-antagonist, naloxone, into the PAG or lesioning of the PAG have blocked the analgesic effects of systemic morphine (Tsou and Jang, 1964; Vigouret et al., 1973; Zhang et al., 1998; Loyd et al., 2008b). These studies highlight the role of the PAG as a crucial site of action for the analgesic effects of locally as well as systemically-administered opioids.

It has been proposed that inhibitory cells within the PAG express the mu-opioid receptor (Osborne et al., 1996; Vaughan et al., 1997), which implies that mu agonists produce analgesia through the disinhibition of PAG output (Vaughan et al., 1997; Budai and Fields, 1998; Heinricher et al., 2009). In support of this model, MOR-expressing PAG neurons have been shown through immunohistochemistry to comprise a subset of all PAG GABAergic neurons, but do not appear to be those that project to the RVM (Reichling and Basbaum, 1990; Kalyuzhny and Wessendorf, 1998; Commons et al., 2000). However, other studies have also shown that PAG neurons containing the mu-opioid receptor comprise over half of RVM-projecting cells (Commons et al., 2000), and these neurons are also presumed to be GABAergic. Thus, the effects of opioid microinjections within the PAG could induce analgesia through inhibition of both local interneurons, RVM-projecting PAG neurons, or both.

Pharmacological and electrophysiological studies have established the clear role of opioidergic circuits in the PAG. However, direct manipulation of PAG circuits has posed challenges due to the molecular complexity of neurons in the PAG. Recent efforts have dissected the role of specific cell types in the PAG using Cre drivers and genetic tools. Using these approaches, the role of PAG neurons in modulating responses to distinct somatosensory modalities have recently been examined in more detail.

Here, we use the MOR^Cre^ allele to determine the function of PAG^MOR^ neurons in the modulation of pain and itch behaviors. We show that activation of PAG^MOR^ neurons generally facilitates nocifensive behaviors but inhibits nociception and responses to pruritogen-evoked itch. These observations may reflect neural circuits that are divergently engaged to modulate nociception and itch depending on behavioral context.

## Results

### Anatomical and neurochemical characterization of PBN neurons projecting to the PAG

Midbrain and brainstem structures, centered upon the PAG, have been shown to modulate nociception (McMullan and Lumb, 2006; Lau and Vaughan, 2014; Wang et al., 2014; Cheriyan and Sheets, 2018). In particular, we focused on inputs to the PAG from the PBN, which have been extensively documented in relaying nociceptive information from the spinal cord to supraspinal targets (Todd, 2010; Cameron et al., 2015). The PBN has also been shown to engage descending pain modulatory systems (Gauriau and Bernard, 2002; Roeder et al., 2016). We have previously shown that stimulating PBN projections within the PAG inhibits reflexive responses in the tail flick assay (Chiang et al., 2020). Our prior observation supports the role of the PBN in the central modulation of pain through a direct connection to the PAG. Building on this work, we wanted to characterize the molecular organization between the PBN and PAG that could represent a descending pathway for the modulation of nociception.

Consistent with our previous findings (Chiang et al., 2020), when we injected the retrograde tracer, cholera toxin B (CTB) into the PAG, we identified extensive labeling of cell bodies within the PBN, supporting the idea that PBN neurons directly project to the PAG (Figure 1A). As an extension of this observation, we sought to neurochemically characterize PBN neurons that project to the PAG using a combination of retrograde tracing and multiplex fluorescent in situ hybridization (FISH). We injected AAVretro-hSyn-eGFP into the PAG of wild type mice and subsequently stained PBN sections containing eYFP and analyzed their expression of known markers of PBN populations that have previously been implicated in modulating pain behaviors including *Tac1, Pdyn*, and *Penk* (Figure 1B) (Han et al., 2015; Barik et al., 2018; Chiang et al., 2020). We identified that among the PBN neurons that project to the PAG, there was considerable overlap with *Tac1* (19.4+/-3.5), but minimal overlap with *Pdyn* (1.6+/-1.6) and *Penk* (1.8+/-1.4) (Figure 1C). These findings suggest that PBN neurons exhibit cell-type specificity with regard to their innervation of the PAG.

**Figure 1.**
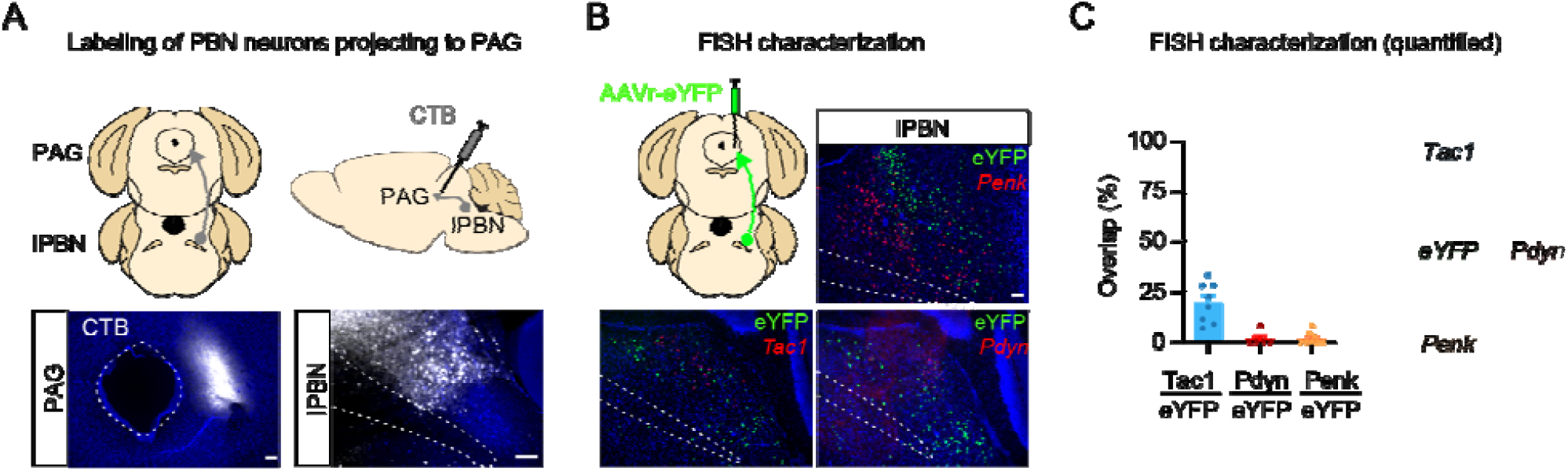
Characterization of PBN neurons projecting to the PAG. (A) Stereotaxic delivery of CTB in the PAG and back labeled cell bodies in the PBN. Representative images are shown. Scale bars = 50 µm. (B) Characterization of PBN neurons that project to the PAG using a retrogradely-labeled reporter (AAVr-eYFP) and FISH. Representative images are shown from 2-3 mice. (C) Percentages of eYFP-expressing PBN neurons expressing Tac1 (19.4+/-3.5), Pdyn (1.6+/-1.6), and Penk (1.8+/-1.4) and their relative overlap are shown. Data are mean + SEM with dots representing 5-8 PBN sections from 2-3 mice. Scale bars = 50 µm.

### Molecular and anatomical characterization of PAG^MOR^ neurons

Immunohistochemical and combined electrophysiological and pharmacological studies have highlighted the existence of neurons containing the mu-opioid receptor within the PAG [36,37,49,50]. For instance, microinjections of mu agonists such as morphine [51] and DAMGO [52] into the PAG inhibit withdrawal reflexes to noxious heat [51] by acting directly on neurons containing the mu-opioid receptor.

The circuitry and molecular identities of mu-sensitive PAG neurons remains unclear. Although it is thought that morphine may inhibit local GABAergic PAG neurons containing the mu-opioid receptor to mediate anti-nociception [50], a significant number of PAG neurons immunoreactive for the mu-opioid receptor have been shown to project directly to the RVM [36]. Furthermore, it also remains unknown whether these opioid-sensitive neurons comprise glutamatergic or GABAergic subpopulations of PAG neurons. We performed tracing analysis of PAG^MOR^ neurons using a Cre-dependent fluorescent reporter (Figure 2A). We identified projection targets in numerous structures throughout the forebrain and brainstem such as the anterior hypothalamus, ventral tegmental area, and RVM (Figure 2A, B).

**Figure 2.**
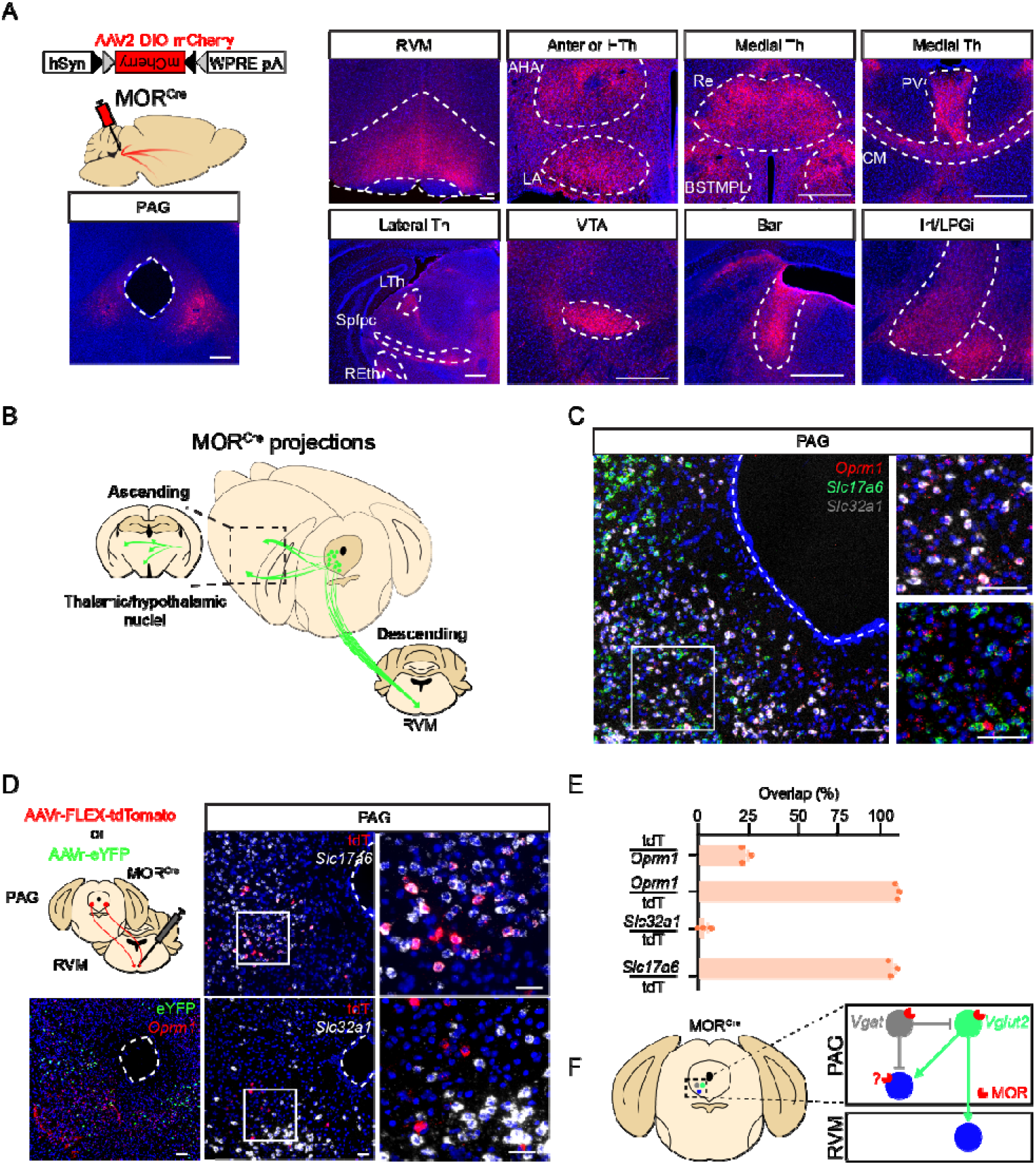
Molecular and anatomic characterization of MOR neurons in the PAG. (A) Approach to visualize projections from PAG^MOR^ neurons using Cre-dependent tracers introduced into the PAG. Major downstream targets of PAG^MOR^ neurons throughout the brain and brainstem. Representative images are shown. Re: reuniens thalamic nucleus, BSTMPL: bed nucleus stria terminalis, medial posterolateral, PV: paraventricular thalamic nucleus, CM: centromedial thalamic nucleus, Spfpc: subparafascicular thalamus, REth: retroethmoid intramedullar thalamic area, PR: prerubral thalamic area, RVM: rostral ventromedial medulla, AHA: anterior hypothalamic area, LA: lateroanterior nuclei, Bar: Barrington’s nucleus, Irt: intermediate reticular nucleus, LPGi: lateral paragigantocellular nucleus, VTA: ventral tegmental area, RRF: retrorubral field. Scale bar = 100 µm. (B) Cartoon representation of PAG^MOR^ ascending and descending projections throughout the brain. (C) FISH characterization of *Oprm1* PAG neurons with respect to excitatory (*Slc17a6)* and inhibitory (*Slc32a1)* markers. Scale bars = 100 µm. (D) Cartoon depiction of two complementary approaches used to characterize descending projections of PAG^MOR^ neurons to the RVM. Retrograde tracers are introduced into the RVM of MOR^Cre^ or WT mice and labeled cell bodies in the PAG are characterized (using *Oprm1, Slc17a6*, and *Slc32a1)* using FISH. Representative images of PAG sections are shown. Scale bar = 50 µm. (E) Quantification of (D). Data are mean + SEM with dots representing individual mice (n=3 mice, with an average of 3-4 sections per mouse). (F) Model for how divergent excitatory and inhibitory PAG neurons containing the mu-opioid receptor (MOR) modulate pain behaviors.

When we stained *Oprm1*-expressing neurons in the PAG, we found that these neurons extensively overlapped with both *Slc32a1 (Vgat)* and *Slc17a6 (Vglut2)* (Figure 2C). This observation supports the idea that PAG^MOR^ neurons comprise a heterogeneous population and that the glutamatergic and GABAergic subpopulations may contribute distinct roles in nociception (Aimone and Gebhart, 1986; Jiang and Behbehani, 2001; Samineni et al., 2017, 2019). Given the important role of the PAG in the descending modulation of pain (Tsou and Jang, 1964; Vigouret et al., 1973; Zhang et al., 1998; Loyd et al., 2008b) and its significant projections to the RVM (Basbaum and Fields, 1979; Gebhart, 1982; Prieto et al., 1983), we focused on this particular pathway. To specifically characterize PAG^MOR^ neurons projecting to the RVM, we injected complementary Cre-dependent and Cre-independent retrograde tracers in the RVM of MOR^Cre^ and wild type mice, respectively (Figure 2D). We then used FISH to characterize back-labeled neurons in the PAG. In contrast to *Oprm1* neurons within the PAG that expressed either *Slc32a1* or *Slc17a6*, PAG^MOR^ neurons that project to the RVM were found to be exclusively glutamatergic by their expression of *Slc17a6* and limited expression of *Slc32a1* (Figure 2E). On the basis of these findings, it is likely that GABAergic PAG^MOR^ neurons participate in the modulation of local circuitry within the PAG whereas glutamatergic PAG^MOR^ neurons project to other structures such as the RVM (Figure 2F).

We then assessed the functional connection between the PAG and RVM using an optogenetic approach in MOR^Cre^ mice (Figure 3A). The optogenetic activation of PAG^MOR^ fibers in the RVM resulted in the robust induction of fos-expressing neurons in animals receiving channelrhodopsin compared to controls (Figure 3B). Behaviorally, photostimulation of PAG^MOR^ fibers within the RVM did not result in heightened jumping responses on the hotplate test that reached statistical significance (Figure 3C). Although stimulation of PAG^MOR^ fibers in the RVM reliably induced activity in the RVM based on fos expression, this stimulation was not sufficient to induce nocifensive behaviors with hotplate testing.

**Figure 3.**
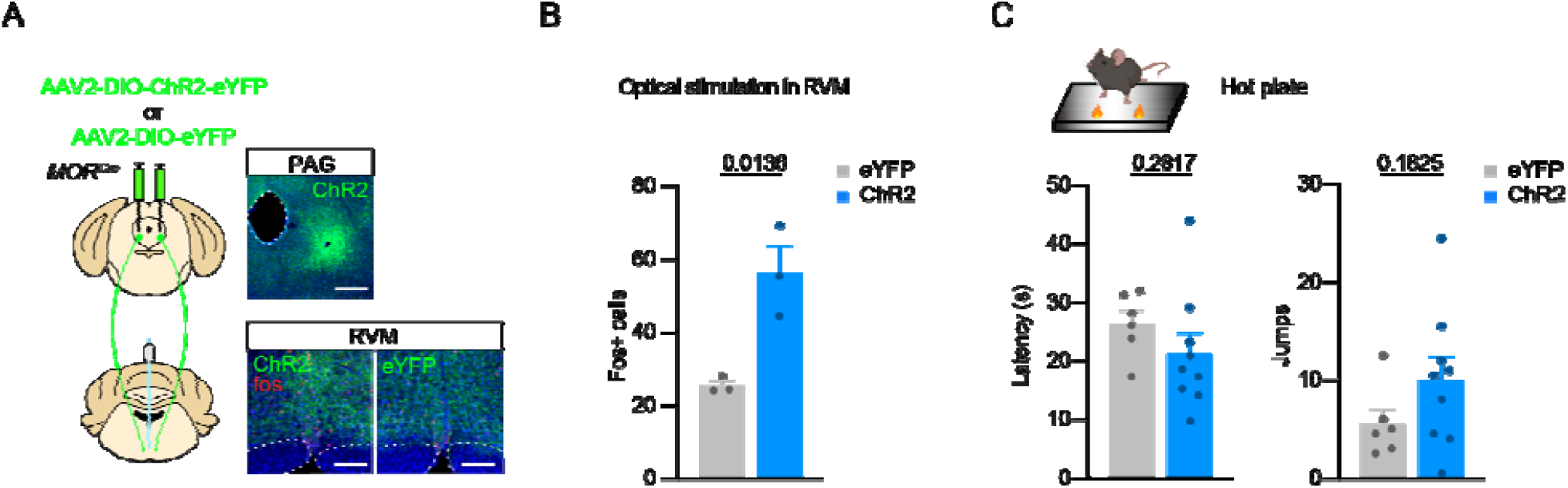
Optogenetic manipulation of PAG^MOR^ projections in the RVM. (A) Approach to optogenetically activate PAG^MOR^ projections to the RVM. The activation of PAG^MOR^ terminals within the RVM induces the expression of fos in the RVM. (B) Quantification of D. Data are mean + SEM with dots representing individual mice (n=3 mice, with an avearge of 3-4 sections per mouse). P value represents the result of two-tailed, unpaired t-test. Representative images from control and ChR2 mice. Scale bars = 200 µm. (C) The effect of optogenetic activation of PAG projections in the RVM on jumping behaviors on the hotplate assay (total jumps and latency to jump). Data are mean + SEM with dots representing individual mice (n=6–9 mice/group). P value represents the result of two-tailed, unpaired t-test.

### PAG^MOR^ neurons modulate complex somatosensory behaviors

Previous studies have shown the dramatic anti-nociceptive effects of pharmacological application of mu agonists within the PAG (Lewis and Gebhart, 1977; Carstens et al., 1990; Yaksh, 1997; Loyd et al., 2008b, 2008a). We therefore sought to test the contributions of PAG^MOR^ neurons by directly targeting them using a chemogentic approach. In contrast to our experiments using optogenetics which did not yield any significant changes in nocifensive behaviors, we reasoned that this approach would allow us to mimic the pharmacological action of infusing mu agonists and antagonists into the PAG. We activated and inhibited PAG^MOR^ neurons by injection of AAV2.hSyn.DIO.hM3Dq-mCherry and AAV2.hSyn.DIO.hM4Di-mCherry, respectively, into the PAG of MOR^Cre^ mice. As a validation of these chemogenetic actuators, the chemogenetic activation of PAG^MOR^ neurons resulted in the induction of fos expression in the PAG (Figure 4A). Consistent with our optogenetic activation of the population, the chemogenetic activation of PAG^MOR^ neurons also activated neurons in the RVM, as determined by fos expression (Figure 4B). Neither activation nor inhibition of PAG^MOR^ neurons affected locomotor activity in an open field assay (Figure 4C), suggesting that the manipulation of PAG^MOR^ neurons does not influence locomotion. To examine the role of PAG^MOR^ neurons in nocifensive behaviors, we tested mice in several noxious pain assays, including hotplate (55° C) and injury models using intraplantar formalin (2% w/v in saline) and capsaicin (using 0.1% w/v capsaicin dissolved in 10% ethanol and saline). On the hotplate assay, the chemogenetic activation of PAG^MOR^ neurons facilitated jumping in naive animals. In contrast, chemogenetic inhibition attenuated jumping in capsaicin-injured animals (Figure 4D). However, neither chemogenetic activation nor inhibition influenced licking behaviors in an acute capsaicin-induced injury model (Figure 4E). To assess the contribution of PAG^MOR^ neurons to a more persistent injury model, we injected mice with intraplantar formalin and quantified their licking behaviors targeted to the injured paw (Figure 4F). We found that activation of PAG^MOR^ neurons modestly enhanced cumulative licking responses over the second phase of the formalin assay, although this trend was not found to be statistically significant (Figure 4G). In contrast, the inhibition of PAG^MOR^ neurons trended toward a reduction in formalin-induced licking, yet this was also not found to be statistically significant. Together, our chemogenetic manipulations of PAG^MOR^ neurons revealed that they robustly modulated jumping behaviors on a noxious hotplate, but did not significantly modulate licking responses in acute injury models.

**Figure 4.**
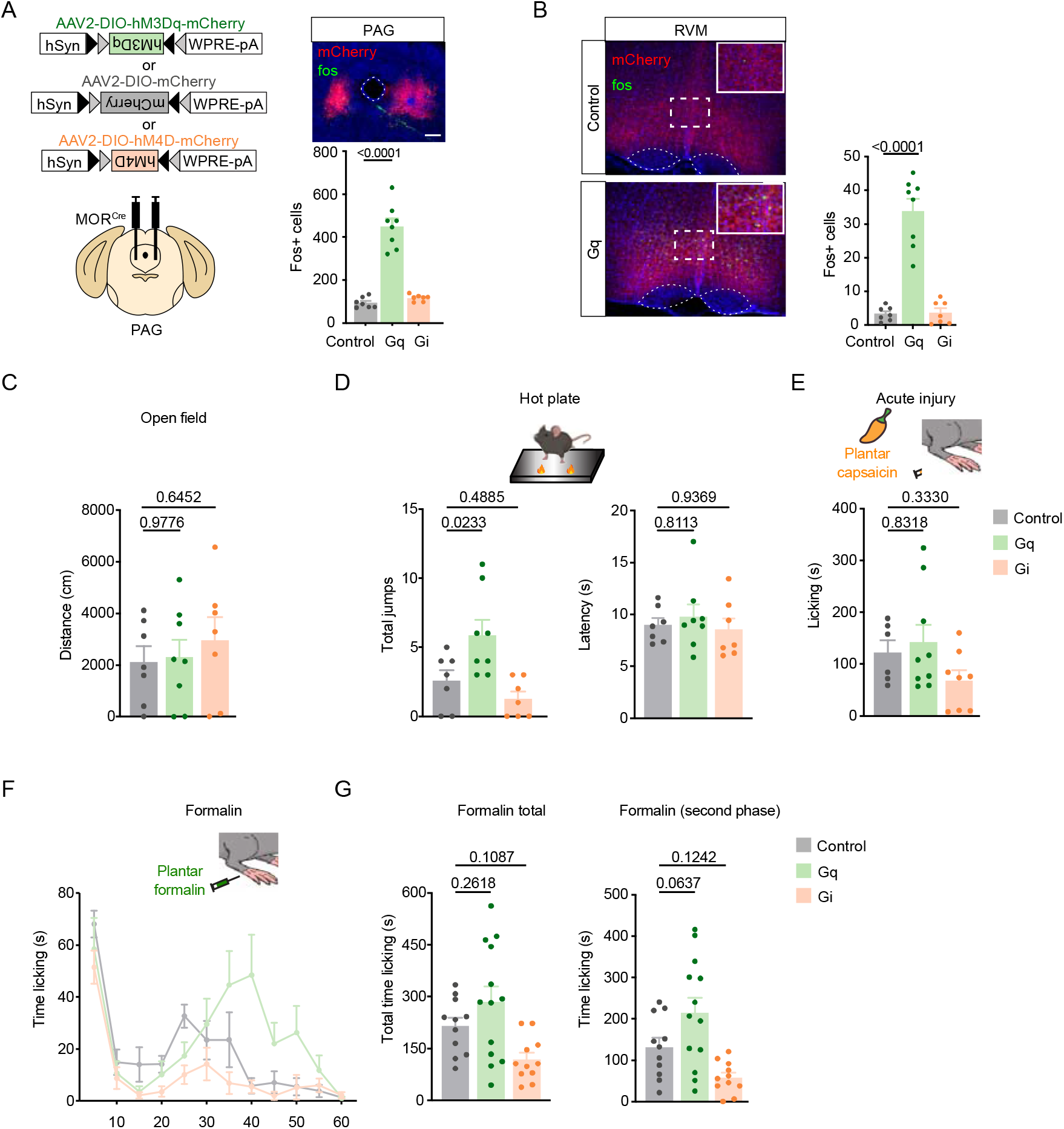
PAG^MOR^ neurons modulate nocifensive behaviors. (A) Approach to chemogenetically activate and inhibit PAG^MOR^ neurons involving the injection of Cre-dependent hM3Dq and hM4Di, respectively to the PAG. A representative image is shown. Scale bar = 50 µm. (B) Induction of fos expression within the RVM following chemogenetic manipulation of PAG^MOR^ neurons. Representative images of RVM sections are shown. Scale bar = 50 µm. Comparison of fos expression in the RVM. Data are mean + SEM with dots representing individual mice (n=7-8 mice per group, averaging 4 sections per mouse). *P* values indicate the results of one-way ANOVA with Dunnett’s multiple corrections test. (C-E) Effect of chemogenetic activation and inhibition on the (C) open field, (D) hotplate test examining total jumps and latency to jump, and (E) nocifensive licking behaviors following intraplantar capsaicin-induced injury. Data are mean + SEM with dots representing individual mice (Control: 6-11, hM3Dq: 7-13, and hM4Di: 7-11 mice per group). *P* values indicate the results of one-way ANOVA with Dunnett’s multiple corrections test. ((F-G) Effect of chemogenetic manipulations of PAG^MOR^ neurons on licking responses following intraplantar formalin. (F) Time course of licking responses binned every 5 min over 1 h. Data are (F) mean +/-SEM and (G) mean + SEM with dots representing individual mice (n=11-13 mice per group. *P* values indicate the results of one-way ANOVA with Dunnett’s multiple corrections test.

Upon assessment of reflexive nociceptive behaviors (Figure 5A), we found that the chemogenetic manipulations of PAG^MOR^ neurons revealed striking complexity in contrast to the behaviors we previously observed on the hotplate assay and with formalin and capsaicin-induced licking. For example, in a model of chloroquine-induced itch, chemogenetic activation of PAG^MOR^ neurons with CNO robustly attenuated scratching responses compared to baseline (Figure 5B). In the von Frey assay, chemogenetic activation and inhibition of PAG^MOR^ neurons bidirectionally modulated mechanical thresholds (Figure 5C). In both the von Frey and Hargreaves assays, activation of PAG^MOR^ neurons consistently inhibited thresholds (Figure 5C, D), in contrast to its facilitation of responses to noxious stimuli, such as hotplate (Figure 4D). Inhibition of PAG^MOR^ neurons also resulted in increased sensitivity to mechanical stimulation. However, tail flick responses were not affected by either chemogenetic activation or inhibition of PAG^MOR^ neurons (Figure 5E).

**Figure 5.**
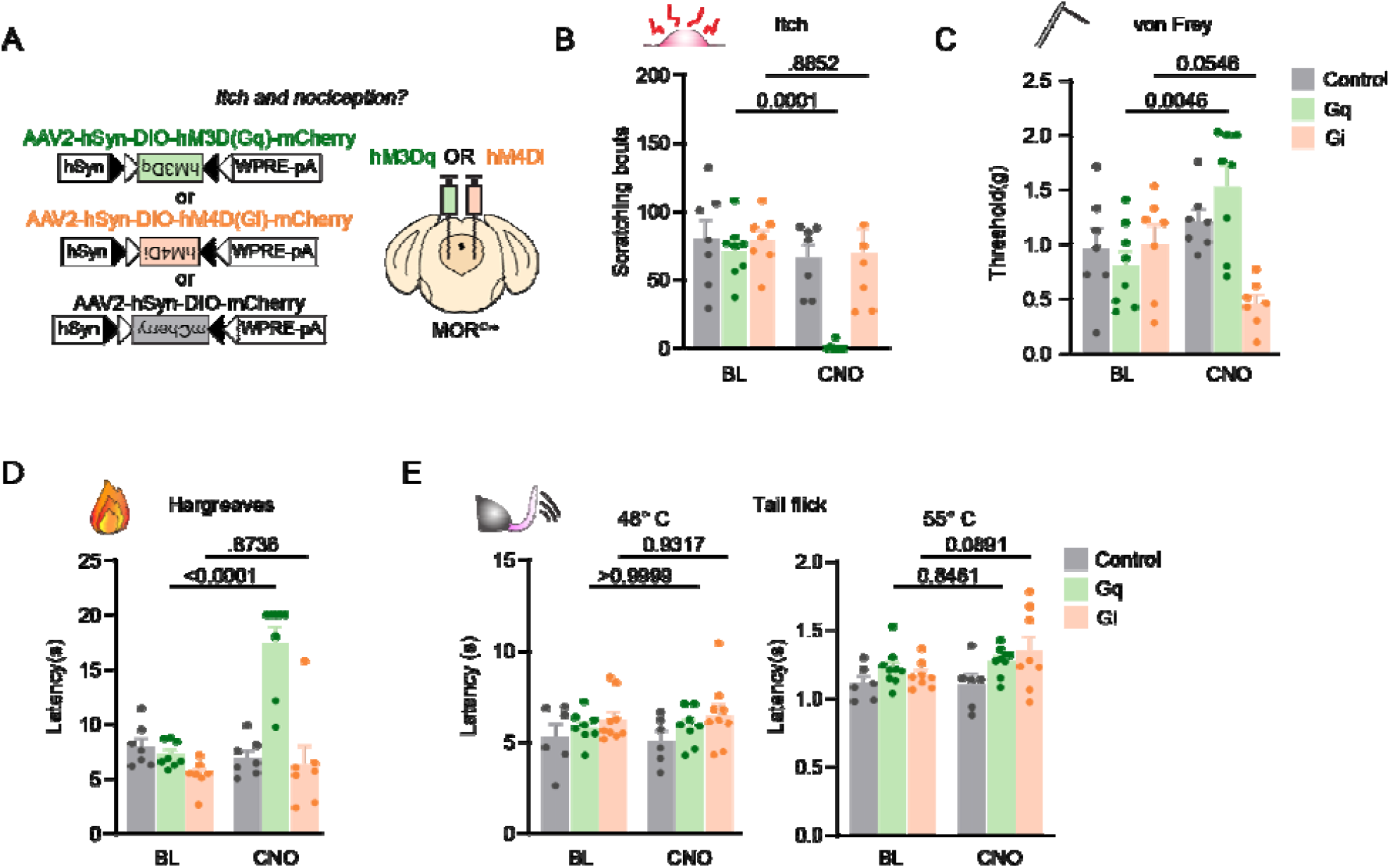
PAG^MOR^ neurons modulate itch and reflexive pain behaviors. ((A) Approach to chemogenetically activate and inhibit PAG^MOR^ neurons involving the injection of Cre-dependent hM3Dq and hM4Di, respectively to the PAG. ((B-E) Effect of chemogenetic manipulations of PAG^MOR^ neurons on (B) chloroquine-induced itch, (C) von Frey assay, (D) Hargreaves assay, and (E) tail flick assay at 48°C and 55° C. Data are mean + SEM with dots representing individual mice (Control: 6-7, hM3Dq: 8-9, and hM4Di: 7-8 mice per group). P values indicate the results of two-way repeated measures ANOVA with Sidak’s multiple corrections test.

Thus, in assays that involved noxious stimulation, activation of PAG^MOR^ neurons generally facilitated nocifensive responses (such as licking and jumping), however for reflexive nociceptive assays such as the von Frey and Hargreaves test, chemogenetic excitation inhibited nociceptive responses. Lastly, the activation of PAG^MOR^ neurons also robustly attenuated chloroquine-induced scratching, representing a divergence from its facilitation of nocifensive responses to hotplate and capsaicin and formalin-induced injury.

### Modulation of latent sensitization by a brainstem circuit

Endogenous opioid signaling has been shown to mask the behavioral expression of pain in a variety of chronic and inflammatory models of pain in a process known as latent sensitization of pain (Li et al., 2001; Le Roy et al., 2011; Corder et al., 2013). Following a noxious injury, animals exhibit a hyperalgesic state that is later suppressed by the upregulation of opioid signaling (Taylor and Corder, 2014). However, the induction of the latent sensitization of pain has been shown to be reversed following the application of opioid receptor antagonists, such as naloxone (Corder et al., 2013). We sought to test the contribution of the PBN to PAG pathway in the latent sensitization of pain (Figure 6A). Consistent with previous findings (Corder et al., 2013), we found that the systemic administration of naloxone (10 mg/kg IP) in naive animals did not affect jumping behaviors (either latency to jump or total jumps) on the hotplate assay compared to saline (Figure B). However, compared to baseline, the administration of naloxone facilitated jumping behaviors in mice that had undergone intracranial surgery (viral injection to the PBN and implantation of optical fibers in the PAG) (Figure 6C). In these mice, naloxone decreased the latency to jump and increased total jumping behaviors (Figure 6C). We found that optogenetic photostimulation of PBN fibers within the PAG blocks the hyperalgesic effects of naloxone and results in the elevation of latencies to jump and reduces total jumps on the hotplate test (Figure 6C). The robust effects of naloxone in precipitating nocifensive behaviors to hotplate testing in the absence of a prior injury (beyond the intracranial viral injection and implant) highlight the role of compensatory endogenous opioids mechanisms that are engaged following cranial surgery. The striking attenuation of naloxone-induced hyperalgesia by optical stimulation of PBN terminals within the PAG suggests that this particular circuit is important for the modulation of opioidergic transmission in latent sensitization.

**Figure 6.**
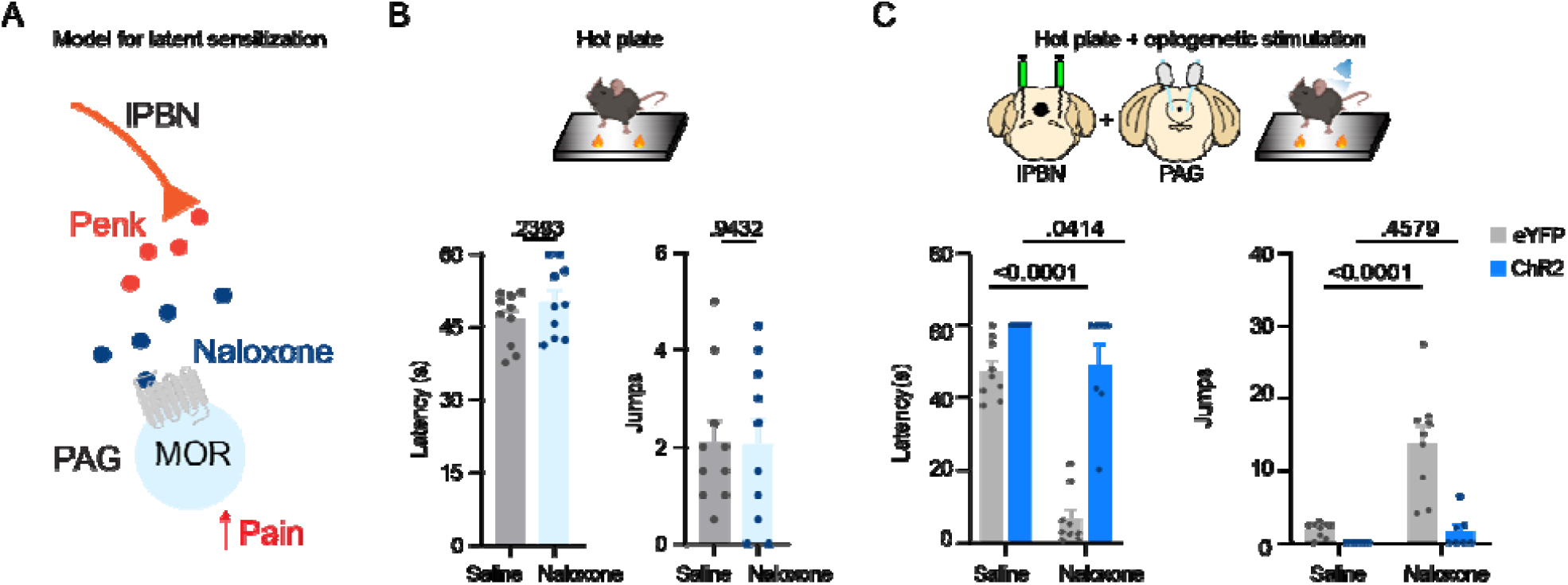
Latent sensitization of pain is modulated by a PBN → PAG pathway. ((A) Model for compensatory endogenous opioid signaling between the PBN and PAG following injury. ((B) Jumping behaviors among naive animals receiving saline or naloxone. Latency to jump and total jumps on a hotplate were quantified. Data are mean + SEM with dots representing individual mice (n=10 mice per group). P values indicate the results of two-tailed unpaired t-test. ((C) Effect of optogenetic stimulation of PBN projections in the PAG in the presence of naloxone on hotplate jumping behaviors including (C) latency to jump and total jumps. Data are mean + SEM with dots representing individual mice (n=7-9 mice per group). *P* values indicate the results of two-way repeated measures ANOVA with Holm-Sidak’s multiple comparisons test.

## Discussion

It has recently been shown that the PAG contains at least two classes of neurons that divergently modulate nociception. In addition to their differences in sensitivity to pharmacological agents (Moreau and Fields, 1986; Carstens et al., 1988; Behbehani et al., 1990), PAG neurons can also be distinguished by their transmitter. GABAergic (Vgat), or inhibitory, neurons are thought to be pain facilitating, whereas glutamatergic (Vglut2), or excitatory, neurons are thought to be anti-nociceptive (Aimone and Gebhart, 1986; Jiang and Behbehani, 2001; Samineni et al., 2017, 2019). In our initial experiments characterizing PAG^MOR^ neurochemically with FISH, we found that *Oprm1* labeled a large number of both glutamatergic and GABAergic neurons within the PAG, which precluded meaningful interpretation of its neurochemical identity given the previous studies that have shown these populations to be differentially engaged for itch and nociception. Therefore, we used a viral approach to label only the PAG^MOR^ neurons that project to the RVM and performed FISH analysis of these neurons. As expected, this approach proved to be specific but not highly sensitive. Although all the neurons back-labeled with tdT were *Oprm1+*, this approach only captured about 25% of the total *Oprm1*-expressing population in the PAG. Therefore, a limitation of this study is that we were only able to determine the molecular identity of a minority of PAG^MOR^ neurons, which may not completely represent all the PAG^MOR^ neurons that could participate in pain-modulatory circuits.

Despite this limitation, we consistently found that RVM-projecting *Oprm1+* neurons in the PAG were glutamatergic and that optogenetic activation of this pathway was sufficient to induce fos expression in the RVM. Our findings suggest that the output of PAG^MOR^ neurons is generally glutamatergic and that the population of gabaergic PAG^MOR^ coule participate in local microcircuits. In this study, we only specifically tested PAG^MOR^ fibers in the RVM using optogenetics on hotplate and did not observe significant results. Additional studies are necessary to determine the contribution of PAG^MOR^ projections to the RVM in other assays. Furthermore, our tracing analysis suggests that perhaps a different circuit, or a combination of circuits, may instead represent the endogenous analgesic pathway centered on PAG^MOR^ neurons.

Another surprising finding in our study was the divergent role of PAG^MOR^ neurons in nocifensive, nociceptive, and itch behaviors. The neurochemical diversity of *Oprm1* may help to explain some of our unexpected behavioral phenotypes. Based on pharmacological studies of mu agonists and antagonists in the PAG, we predicted that the optogenetic or chemogenetic activation of PAG^MOR^ neurons would have facilitated nociception whereas chemogenetic inhibition of these neurons would have anti-nociceptive effects. We found that this paradigm was generally predictive when it came to noxious stimuli or inflammatory and acute injury models. With a 55° C hotplate, optogenetic and chemogenetic activation both elicited jumping behaviors. We saw a similar trend with intraplantar capsaicin and formalin, although these trends did not reach statistical significance. Intriguingly, however, we generally observed the opposite pattern with assays more commonly used to assess nociceptive thresholds. In contrast to the hotplate, formalin, and capsaicin assays, the von Frey and hargreaves assays revealed that activation and inhibition of PAG^MOR^ neurons were anti- and pro-nociceptive, respectively.

One possible explanation for our divergent observations with nocifensive compared to nociceptive testing is that formalin, capsaicin, and a 55° C hotplate may have led to induction of an injury state that exposed the pain-facilitating role of PAG^MOR^ neurons, which is not observed in non-noxious assays for nociception. How these circuits are differentially engaged by the stimulus is an important question. We have recently shown that a population of RVM neurons that participate in both the inhibition of pain and itch are also engaged during stress (Nguyen et al., 2022). It is likely that stress also contributes to the generally anti-nociceptive effects of chemogenetic activation of PAG^MOR^ neurons in the context of hotplate, formalin, and capsaicin-induced injury as all of these assays involve more handling than hargreaves and von Frey testing. The descending modulatory system has been shown to be important in the switch from acute stress-induced analgesia and chronic stress-induced hyperalgesia (François et al., 2017). Therefore, further studies are necessary to determine the specific role of PAG neurons in coordinating specific behavioral states, particularly in the context of acute and chronic stress.

Another intriguing finding was that activation of PAG^MOR^ neurons completely attenuated chloroquine-induced itch. This observation that PAG neurons are dichotomously engaged for nociception and pruriception would align with recent work showing that both GABAergic and glutamatergic PAG neurons divergently modulate itch and pain (Samineni et al., 2017, 2019). Our exploration into the function of PAG circuitry using PAG^MOR^ neurons as an entry-point supports the previous observation that neurons in the PAG do not participate in all forms of nociception and pruriception homogeneously.

Lastly, we found that activation of PBN projections in the PAG reversed naloxone-induced hyperalgesia in a potential new model of latent sensitization. It is notable that we did not perform an injury model as is classically done in assessing latent sensitization (such as with plantar incision, nerve injury, or complete Freund’s adjuvant (CFA)) (Corder et al., 2013; Marvizon et al., 2015). Instead, the administration of naloxone in animals that had previously received cranial implants of optical fibers and stereotactic viral injections was sufficient for the emergence of hyperalgesia. Our results suggest that intracranial surgeries are sufficient to induce compensatory analgesia and this may represent a contributor to chronic pain in mice. This presents an important consideration for future studies using stereotactic delivery or intracranial implantation as these manipulations will likely affect endogenous mu-opioid receptor tone.

## Methods

### Mice

All animals were of the C57BL/6J background. MOR^Cre^ mice were a generous gift from Richard Palmiter and are now available on Jax #034475. The studies were performed in both male and female mice 8-10 weeks of age. Even numbers of male and female mice were used for all experiments and no clear sex differences were observed so data were pooled. Mice were given free access to food and water and housed under standard laboratory conditions. The use of animals was approved by the Institutional Animal Care and Use Committee of the University of Pittsburgh.

### Viral vectors

All viruses are commercially available from UNC Vector Core and Addgene. AAV2.hSyn.DIO.hM4D(Gi)-mCherry, AAV2-hsyn-DIO-mCherry, AAV2-hsyn-DIO-hM3D(Gq)-mCherry, AAV2/EF1a-DIO-hChR2(H134R)-eYFP, AAV2.EF1a.DIO.eYFP, pENN.AAV.hSyn.HI.eGFP-Cre.WPRE.SV40, AAVr-Ef1a-eYFP, and AAVr-hsyn-DIO-mCherry.

### Stereotaxic injections and optical fiber implantation

Animals were anesthetized with 2% isoflurane and placed in a stereotaxic head frame. A drill bit (MA Ford, #87) was used to create a burr hole and a custom-made metal needle (33 gauge) loaded with virus was subsequently inserted through the hole to the injection site. Virus was infused at a rate of 100 nL/min using a Hamilton syringe with a microsyringe pump (World Precision Instruments). Mice received 250-500 nL of virus. The injection needle was left in place for an additional 5 min and then slowly withdrawn over another 5 min. Injections and cannula implantations were performed at the following coordinates for each brain region: RVM: AP −5.80 mm, ML 0.00 mm, DV −6.00, PAG: AP −4.70 mm, ML ± 0.74 mm, DV: −2.75 at a 4° angle, and

PBN: AP −5.11 mm, ML ± 1.25 mm, DV: −3.25. For implantation of optical fibers (Thor Labs: 1.25 mm ceramic ferrule 230 mm diameter), implants were slowly lowered 0.3 - 0.5 mm above the site of injection and secured to the skull with a thin layer of Vetbond (3M) and dental cement.

The incision was closed using Vetbond, and animals were provided analgesics (ketofen, i.p. 10 mg/kg; buprenorphine, subcutaneous, 0.3 mg/kg) and allowed to recover over a heat pad. Mice were given 4 weeks to recover prior to experimentation.

### Pharmacologic agents

Clozapine-N-oxide (Tocris) was dissolved in PBS and administered intraperitoneally (5 mg/kg). Capsaicin (0.1%) in 10% EtOH in PBS was injected 10 µL into the plantar hindpaw. Chloroquine diphosphate salt (Sigma) was dissolved in physiological saline (100 µg in 10 µL) and administered intradermally. Formalin (Sigma; 2% w/v) in saline was injected 10 µL into the plantar hindpaw.

### RNAscope in situ hybridization

Multiplex fluorescent in situ hybridization (FISH) was performed according to the manufacturer’s instructions (Advanced Cell Diagnostics #320850). Briefly, 14 µm-thick fresh-frozen sections containing the RVM were fixed in 4% paraformaldehyde, dehydrated, treated with protease for 15 min, and hybridized with gene-specific probes to mouse. Probes were used to detect Probe-eYFP-C3 Cat No. 312131-C3, Mm-Penk-C1 (#318761), tdTomato-C2 (#317041-C2), mCherry-C2 (#431201), Mm-Oprm1-C1 (#315841), Mm-Tac1-C1 (#517971), Mm-Pdyn-C2 (#31877), Mm-Slc32a1-C3 (#319191), and Mm-Slc17a6-C3 (#319171). DAPI (#320858) was used to visualize nuclei. 3-plex positive (#320881) and negative (#320871) control probes were tested. Three to four z-stacked sections were quantified for a given mouse, and 3-4 mice were used per experiment.

### Immunohistochemistry

Mice were anesthetized with an intraperitoneal injection of urethane, transcardially perfused, and post-fixed at least 4 h in 4% paraformaldehyde. 40 µm thick brain sections were collected on a cryostat for immunohistochemistry. Sections were blocked at room temperature for 2 h in 5% donkey serum, 0.2% triton, in phosphate buffered saline. Primary antisera was incubated for 14 h overnight at 4° C: rabbit anti-RFP (1:1K), chicken anti-GFP (1:1K), and mouse anti-NeuN (1:500). Sections were subsequently washed three times for 20 min in wash buffer (0.2% triton, in PBS) and incubated in secondary antibodies (Life Technologies, 1:500) at room temperature for 2 h. Sections were then incubated in Hoechst (ThermoFisher, 1:10K) for 1 min and washed 3 times for 15 min in wash buffer, mounted and cover slipped.

### Fos experiments

Mice received either optical stimulation or CNO as described for behavioral testing. Brain tissues were harvested 90 min after for immunohistochemistry. For optogenetically-induced fos expression, mice were photostimulated for 20 min at a 3 s on, 2 s off stimulation pattern and subsequently perfused 90 min after the initial onset of photostimulation as noted for immunohistochemistry.

### Image acquisition and quantification

Full-tissue thickness sections were imaged using either an Olympus BX53 fluorescent microscope with UPlanSApo 4x, 10x, or 20x objectives or a Nikon A1R confocal microscope with 20X or 60X objectives. All images were quantified and analyzed using ImageJ. To quantify images in RNAscope in situ hybridization experiments, confocal images of tissue samples (3-4 sections per mouse over 3-4 mice) were imaged and only cells whose nuclei were clearly visible by DAPI staining and exhibited fluorescent signal were counted.

### Behavior

All assays were performed in the Pittsburgh Phenotyping Core and scored by an experimenter blind to treatment. For all chemogenetic behavioral experiments, CNO was administered (5 mg/kg IP) 30 min prior to the start of behavioral testing. Mice that were optogenetically tested were photostimulated with the following parameters: 10 mW laser power, 20 Hz stimulation frequency, and 5 ms pulse duration.

### Observation of scratching behavior

Scratching behavior was observed using a previously reported method (Kardon et al., 2014). On the testing day, the mice were individually placed in the observation cage (12×9×14 cm) to permit acclimation for 30 min. Scratching behavior was videotaped for 30 min after administration of chloroquine. The temporal and total numbers of scratch bouts by the hind paws at various body sites during this period were counted. For the experiments targeting the lumbar spinal cord, the amount of time spent biting following intradermal chloroquine was summated.

### Hargreaves testing

Animals were acclimated on a glass plate held at 30° C (Model 390 Series 8, IITC Life Science Inc.). A radiant heat source was applied to the hindpaw and latency to paw withdrawal was recorded (Hargreaves et al., 1988). Two trials were conducted on each paw, with at least 5 min between testing the opposite paw and at least 10 min between testing the same paw. To avoid tissue damage, a cut off latency of 20 s was set. Values from both paws were averaged to determine withdrawal latency.

### Tail-flick assay

Mice were placed in custom-made mouse restraints and allowed to habituate 15 min before testing. Tails were immersed 3 cm into a water bath at 48° and 55° C, and the latency to tail-flick was measured three times with a 1 min interval between trials. For optogenetic testing, mice were photostimulated for 10 s prior to testing. Cut-offs times were implemented at 25 s (48°) and 5 s (55° C) to prevent tissue damage.

### Hotplate

Mice were placed on a 55° C hotplate, and the latency to first nocifensive response (jump) and total number of jumps were measured over a 60 s period. Values were averaged across two trials for each mouse spaced several minutes apart. For optogenetic testing, mice were photostimulated for 10 s prior to testing.

### Open field

Spontaneous activity in the open field was conducted over 30 min in an automated Versamax Legacy open field apparatus for mice (Omnitech Electronics Incorporated, Columbus, OH). Distance traveled was measured by infrared photobeams located around the perimeter of the arenas interfaced to a computer running Fusion v.6 for Versamax software (Omnitech Electronics Incorporated) which monitored the location and activity of the mouse during testing. Activity plots were generated using the Fusion Locomotor Activity Plotter analyses module (Omnitech Electronics Incorporated). Mice were placed into the open field 30 min post CNO injection.

### Acute injury models

For formalin-induced injury, 10 µL of 2% formalin was injected into the intraplantar hindpaw. Mice were video recorded for 1 hr after formalin injection and time spent licking and lifting the paw was scored in 5-min bins. For capsaicin-induced injury, animals received 10 µL of intraplantar capsaicin (0.1% w/v in 10% ethanol diluted in saline) and the total time spent licking the injured paw was quantified over 20 min. Hargreaves and von Frey testing following capsaicin-induced injury occurred 20 min and 1 h after intraplantar injection.

### Statistical analysis

Statistical analyses were performed using GraphPad Prism 8. Values are presented as mean +/-SEM. Statistical significance was assessed using tests indicated in applicable figure legends. Significance was indicated by p < 0.05. Sample sizes were based on pilot data and are similar to those typically used in the field.

## Acknowledgements

Research reported in this publication was supported by the Virginia Kaufman Endowment Fund, NIH/NIAMS grant AR063772, NIH/NINDS grant NS096705 (S.E.R.), NRSA F31 grant F31NS113371 and NIGM/NIH T32GM008208 (E.N.).

## Author contributions

Conceptualization: E.N., M.C.C., and S.E.R.

Methodology: E.N., M.C.C., and S.E.R.

Investigation: E.N., M.C.C., and C.N.

Writing: E.N. and S.E.R.with input from all authors

Funding Acquisition: E.N. and S.E.R.

